# Tissue Tension and Strain as Indicators of Suction-mediated Cutaneous DNA Transfection: A Parametric Study

**DOI:** 10.1101/2024.02.15.580412

**Authors:** Nandita C. Jhumur, Emran O. Lallow, Catherine Nachtigal, Dahee Yeo, Ijoo Kwon, Young K. Park, Christine C. Roberts, Jeffrey D. Zahn, David I. Shreiber, Jerry W. Shan, Jonathan P. Singer, Joel N. Maslow, Hao Lin

## Abstract

Cutaneous suction-based transfection is a recently developed technique that is painless and simple-to-use for the delivery of DNA for nucleic-acid-based vaccines. The technique promises high efficiency for both antigen expression and immunogenicity as demonstrated in both animal studies and human clinical trials. To realize this promise, a parametric study and systematic evaluation on the efficacy of cutaneous suction as a transfection method was performed. Using Green Fluorescent Protein (GFP) plasmid expression as a transfection reporter in a rat model, the expression level as a function of both suction nozzle size and suction pressure was quantified. A numerical model was employed to compute skin deformation in terms of strain, which was used to correlate with GFP expression. Based on these results, two quantities, total integrated strain and tissue tension, are proposed as indicators of expression level that can be used to guide protocol development and optimization. These indicators are also discussed in relation to possible cellular uptake mechanisms.

## Introduction

In vivo transfection is the introduction of nucleic acids (DNA or RNA) into cells of living organisms to express the encoded proteins. It has broad applications including protein replacement therapy, nucleic-acid-based vaccination, gene therapy for neurodegenerative disorders, hypertension and cardiovascular diseases treatment, cancer treatment, AAV (adeno-associated virus) cloning for clinical therapeutics, cutaneous wound repair, and AeTEP-1 transfection for reducing dengue infection, among many others.^[1-8]^ Transfection platforms employ all of chemical, biological, and physical methods.^[9-13]^ For example, delivery of mRNA encapsulated into lipid nanoparticles belongs to the category of chemical method.^[14]^ DNA delivery more typically employs biological or physical methods, such as the use of viral vectors, electroporation (EP), gene guns, DNA tattooing, sonoporation, optical transfection, magnetofection, or hydrodynamic delivery, with EP emerging as the most commonly used in recent years.^[15-31]^ When compared with chemical and biological delivery modalities, physical transfection methods are attractive due to the avoidance of potential adverse effects associated with chemical or biological agents. On the other hand, they encounter other limitations and potential side effects such as cell/tissue damage or tissue edema.^[32]^

In our prior work, we have demonstrated successful cutaneous, suction-based DNA transfection in both laboratory and clinical applications.^[13]^ Cutaneous DNA administration presents a simple method of cargo delivery, with the additional benefits of access to large numbers of sentinel cells such as keratinocytes and antigen-presenting cells, which are beneficial for vaccination.^[16]^ In our technique, a negative pressure is applied to the skin surface atop the site of shallow DNA injection (Fig. 1). In a rat model, this method induced efficient transfection when compared with DNA injection alone. Moreover, subsequent studies demonstrated that, when benchmarked against jet microdroplet delivery and EP, suction-mediated transfection of the GLS-5310 COVID-19 vaccine induced similar B cell responses but greater T cell responses.^[33]^ These promising results enabled rapid translation of a suction-mediated delivery device into two DNA vaccine clinical trials (NCT04673149; NCT05182567). Clinical trial results confirmed our preclinical findings: suction post-DNA vaccine injection produced equivalent B cell and superior T cell responses in humans when benchmarked against all currently deployed COVID vaccines.^[34,35]^ Importantly for patients, the suction technique caused no pain or trauma to the skin or muscle, unlike what is experienced by other modalities such as EP or air-jet delivery, and the use of naked plasmids reduces toxicity and avoids possible adverse effects in chemical transfection, e.g., systematic inflammatory responses, hepatotoxicity caused by the LNPs in mRNA vaccines, or PEG sensitivity when using PEG as an adjuvant.^[34,36-40]^ Finally, this technique is cost-effective and highly intuitive to use, requiring minimal to no training. Building upon the rapid development and translation into clinical phase studies, this emerging technology has potential to continue to improve and optimize beyond the initial success. In a follow-up work, we have examined the spatial distribution for variable-sized cargoes (dextran, nanoparticles, and DNA) within an intradermal injection, which is an important aspect for secondary delivery mechanisms as their action sites need to co-localize with the cargo.^[41]^

**Figure 1:**
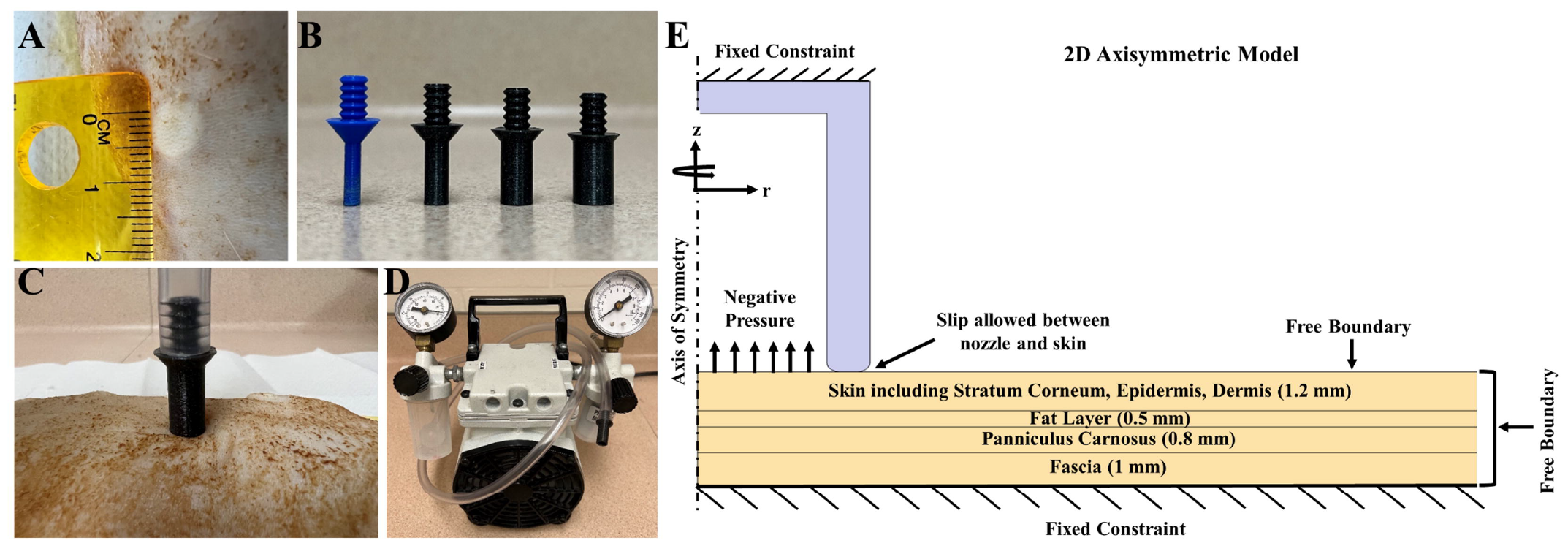
Suction application. (A) Post-injection bleb on rat skin. (B) 3D printed suction nozzles with variable inner diameters (2, 4, 6, and 8 mm). (C) Suction application on rat skin with a suction nozzle. (D) Vacuum pump connects to the suction nozzle through a plastic tube. (E) A schematic of the tissue model.

In the current work, we continue the effort and perform a systematic study of design elements of suction-based cutaneous transfection. Using numerical simulation to correlate expression with the magnitude of tissue deformation, we find that two indicators can predict transfection efficiency: total integrated strain and tissue tension. These indicators can be conveniently and rapidly evaluated to guide parametric study and optimization without resorting to the trial-and-error experiments. We also discuss the results in relation to possible cellular mechanisms for transfection.

## Materials and Methods

### Animals

Male Sprague-Dawley (NTac-SD) rats, 7-11 weeks old, with murine pathogen-free standard from Taconic Biosciences (Germantown, NY) were used. All the rats were kept in controlled environments maintaining 12-hour light: 12-hour dark cycles along with other guidelines per Rutgers University Institutional Animal Care and Use Committee (IACUC).

### GFP Plasmid

pEGFP-N1 DNA plasmid was provided by GeneOne Life Science (Seoul, South Korea). The plasmid was diluted with 1× PBS to a final concentration of 0.5 mg/ml for all the experiments. 50 μL of the solution was used per injection with a total mass of 25 μg DNA plasmid. A 28G insulin syringe (BD, Franklin Lakes, NJ) was used to deliver the solution into rat skin intradermally using a Mantoux injection technique.

### Suction Application

The rats were maintained under anesthesia using Isoflurane, USP Inhalation Anesthetic (Dechra, Overland Park, KS) during the experiments following IACUC protocol 201800077. The rat skin was first shaved with a hair clipper (WAHL, Sterling, Illinois), and a depilatory cream was applied (Nair, Church & Dwight Co., Inc., Ewing, NJ). After 5 minutes, the skin was thoroughly cleaned with 70% ethanol. The Mantoux technique was used for the ID injections targeting the epidermis layer of skin. The diameter of each injection bleb was measured using a ruler (Fig. 1A). The suction nozzles were designed in CAD (SOLIDWORKS, Waltham, MA) and 3D printed with PLA plastic (Fig. 1B). The rim width of each nozzle was kept at 0.8 mm. The suction nozzles were vertically placed on the injection blebs (Fig. 1C), and the threads at the base of the nozzles were fitted to the tube of a vacuum pump (Welch, Mt Prospect, IL; Fig. 1D). Suction pressure was applied for 30 seconds. After the experiments, all animals were single housed. Animals were euthanized via CO_2_ asphyxiation 24-hours after transfection.

### Imaging and Quantification

Skin was excised at the location of injections and placed on glass slides (Fisher Scientific, Pittsburgh, PA) for imaging. Fluorescence microscope (OLYMPUS IX80 and IX81, Olympus, Tokyo, Japan) was used to take images of the skin samples using a 4× objective through FITC and TRITC channels. Exposure settings were kept constant for all images. The images were then processed and quantified with a custom-programmed MATLAB algorithm (R2022a, MathWorks, Natick, MA). The TRITC images were considered representative of background fluorescence and were used to remove them from corresponding FITC images. The Total Pixel Count was calculated as the total number of positive (non-zero and non-negative) pixels after background subtraction, and is akin to the expression area as viewed from the top of the skin surface. The Total Intensity of an image was calculated as the sum of the intensities of all of the positive (non-zero and non-negative) pixels. A standardized positive control value was established for Total Pixel Count and Total Intensity respectively by quantifying expression levels with DNA injection only (same dosage and volume) on each rat without ensuing suction, and then averaging across the animals. Normalized values were then generated by dividing Total Intensity or Total Pixel Count by the respective standardized control value. These values represent fold-enhancement induced by suction when compared with DNA injection alone.

### Simulation

COMSOL Multiphysics 5.5 (Burlington, MA) was used to build a 2D axisymmetric model including skin, fat, panniculus carnosus, and fascia (Fig. 1E) following prior work.^[13]^ The thickness of the layers was determined from histopathological images of rat skin tissue. The mechanical properties of the layers were collected from the literature. An axisymmetric cup was also included. A negative pressure was applied to the top surface of the tissue within the boundary of the cup. The cup was not fixed to the skin, and slip was allowed between the cup and the tissue surface. The strain magnitude was computed as the Frobenius norm of the strain tensor, **E**,

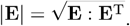

More details, including the hyperelastic energy functions and properties used for each layer and model parameters, can be found in our prior work.^[13]^

### Statistics

Statistical analysis was performed with GraphPad Prism (10.1.2, GraphPad Software, San Diego, CA) using one-way ANOVA with Tukey’s comparisons, where data significance is represented as *P ≤ 0.05, **P ≤ 0.01, ***P ≤ 0.001,and ****P ≤ 0.0001. Data are presented as mean ± SD.

## Results

### Suction applied post-DNA-injection, but not pre-injection, enhances transfection

We first investigated whether the order of injection and suction application affects transfection. For all cases, the injection was 25 μg of pEGFP-N1 in 50 μL of 1× PBS solution; the suction pressure was 80 kPa and applied for 30 seconds with a 6-mm-diameter nozzle. As demonstrated in Figure 2, expression for suction before injection was lower than the injection-only controls, whereas expression increased for suction after injection.

**Figure 2:**
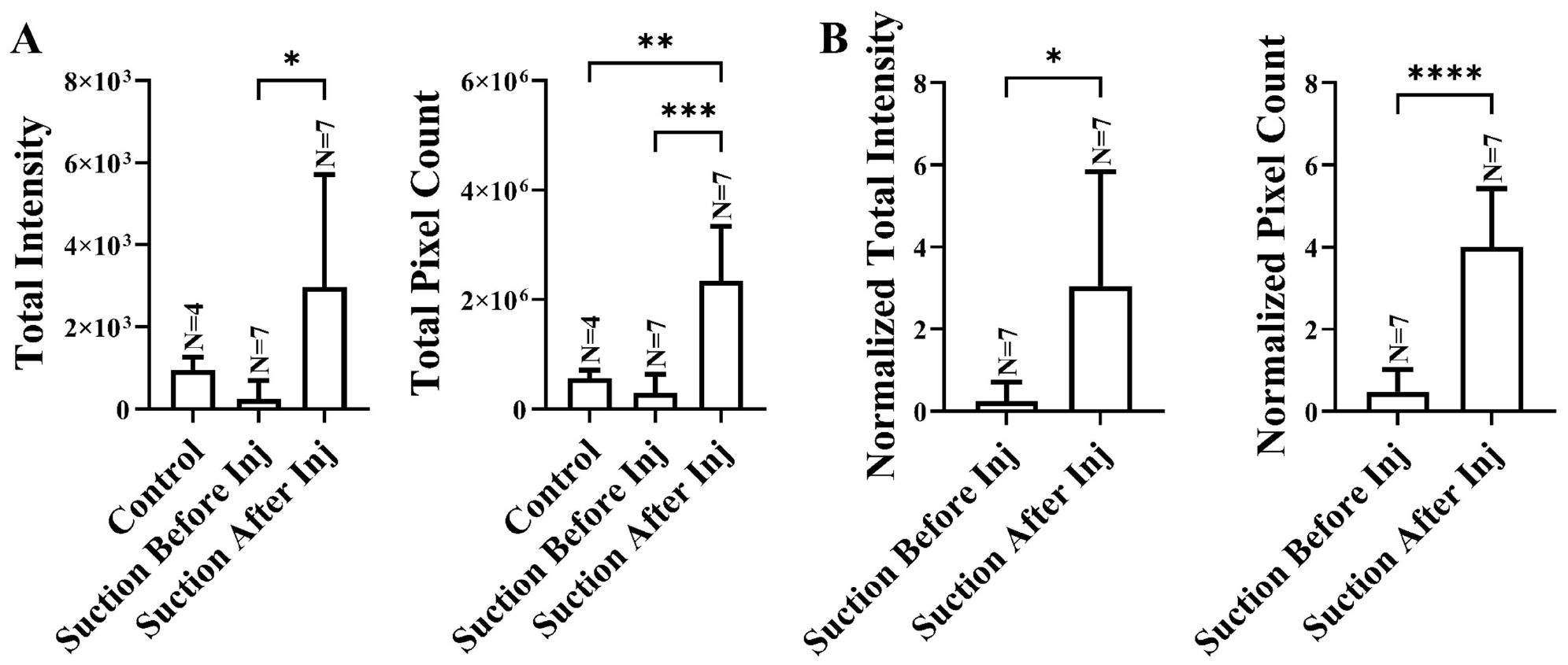
Comparison of the effect of suction application before and after pEGFP-Nl injection on GFP expression after 24 hours. (A) Total intensity (mean+SD) and total pixel count (mean+SD) of GFP expression. (B) Normalized intensity (mean+SD) and normalized pixel count (mean+SD) of GFP expression. 80 kPa suction pressure was applied with a 6 mm nozzle for 30 seconds.

### Expression area and pattern is dependent on nozzle size

Next, we performed a parametric study of the effects of nozzle-size and pressure on suction-mediated transfection. All treatments were performed with suction immediately following ID injection of 25 μg of pEGFP-N1 in 50 μL of 1× PBS solution. Four different circular nozzle opening diameters (2, 4, 6, and 8 mm) were used (Figs. 1B and 3A inset). Figure 3A shows bright field images of the rat skin surface post-injection and suction with the various nozzle sizes. Images are noted to indicate the change in injection bleb site with respect to nozzle size. For 2– and 4-mm openings, the opening rim was within the bleb; the 6-mm rim nearly exactly coincided with the bleb periphery, whereas the 8-mm rim fully enclosed the bleb. These observations have important implications, which we discuss below.

**Figure 3:**
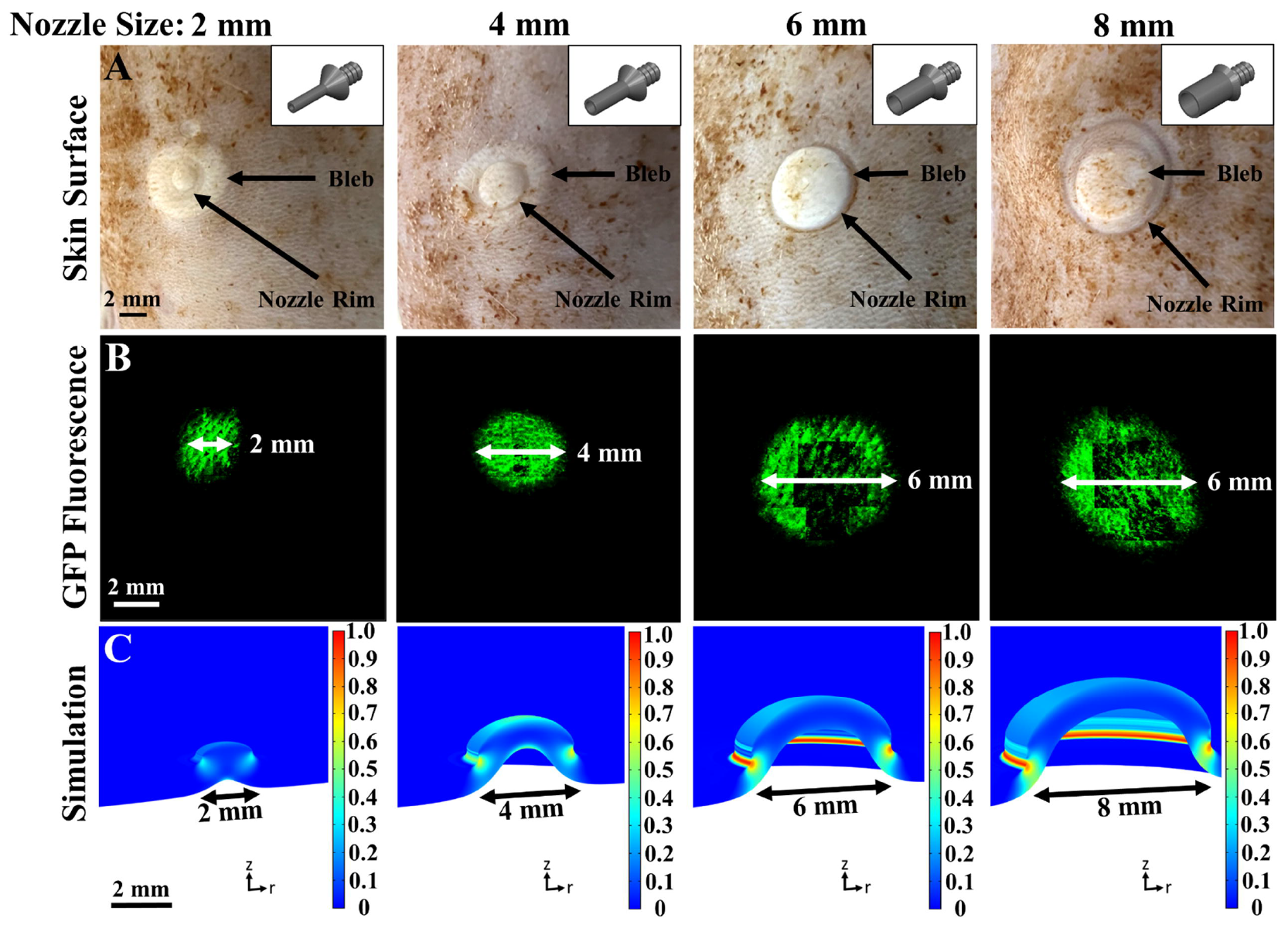
Effect of nozzle size on cutaneous transfection. (A) Injection bleb and nozzle rim impression. 50 μL pEGFP-Nl was ID-injected, followed by application of 65 kPa suction pressure for 30 seconds. (B) 4x objective top-view images of GFP expression in rat skin 24 hours post-suction. (C) 3D view of skin deformation and color-mapped strain following simulation with 65 kPa negative pressure.

Figure 3B shows images of GFP expression in the dissected rat skin 24 hours after treatment. The GFP expression areas were approximately circular in shape in accordance with the circular nozzles. The area of expression is also clearly correlated with the nozzle size. However, it is peculiar to note the change in fluorescence distribution: for the 2– and 4-mm openings, expression was more uniform within the nozzle-covered area; for 6– and 8-mm openings, expression was greater towards the rim. Additionally, for the 2-, 4-, and 6-mm cases, GFP expression was spread within the entire nozzle-covered area reaching the rim; for the 8-mm case, where the nozzle rim was outside of the injection bleb, expression was similar to the bleb size. This result is consistent with previous observations demonstrating that DNA cargo typically remains within the bleb area.^[41]^

### Spatial expression pattern is correlated to strain pattern

Results of the numerical simulations demonstrate a correlation between the tissue strain and GFP expression patterns. Figure 3C shows three-dimensional images of COMSOL simulation of negative-pressure application on the tissue model surface. The skin deformation and strain magnitude (color) for different nozzle sizes using a surface load of 65 kPa negative pressure are shown. For 2– and 4-mm openings, although the strain is concentrated at the suction rim, its magnitude at that location is approximately the same as within the aspirated tissue dome. For larger openings, strain becomes dominantly stronger towards the rim. These strain concentration patterns are echoed in the top-view of GFP expression patterns.

### Expression strength depends on both suction pressure and nozzle size

We performed experiments to systematically quantify the dependence of expression on both suction pressure and nozzle size. For each nozzle size, we applied 35, 50, 65, or 80 kPa of suction pressure for 30 seconds. For the 6-mm nozzle, which was used in clinical trials,^[33,34]^ we added pressures of 70, 75, 85, and 90 kPa. Several general trends are observed in Fig. 4, for both raw or normalized intensity and pixel counts.

**Figure 4:**
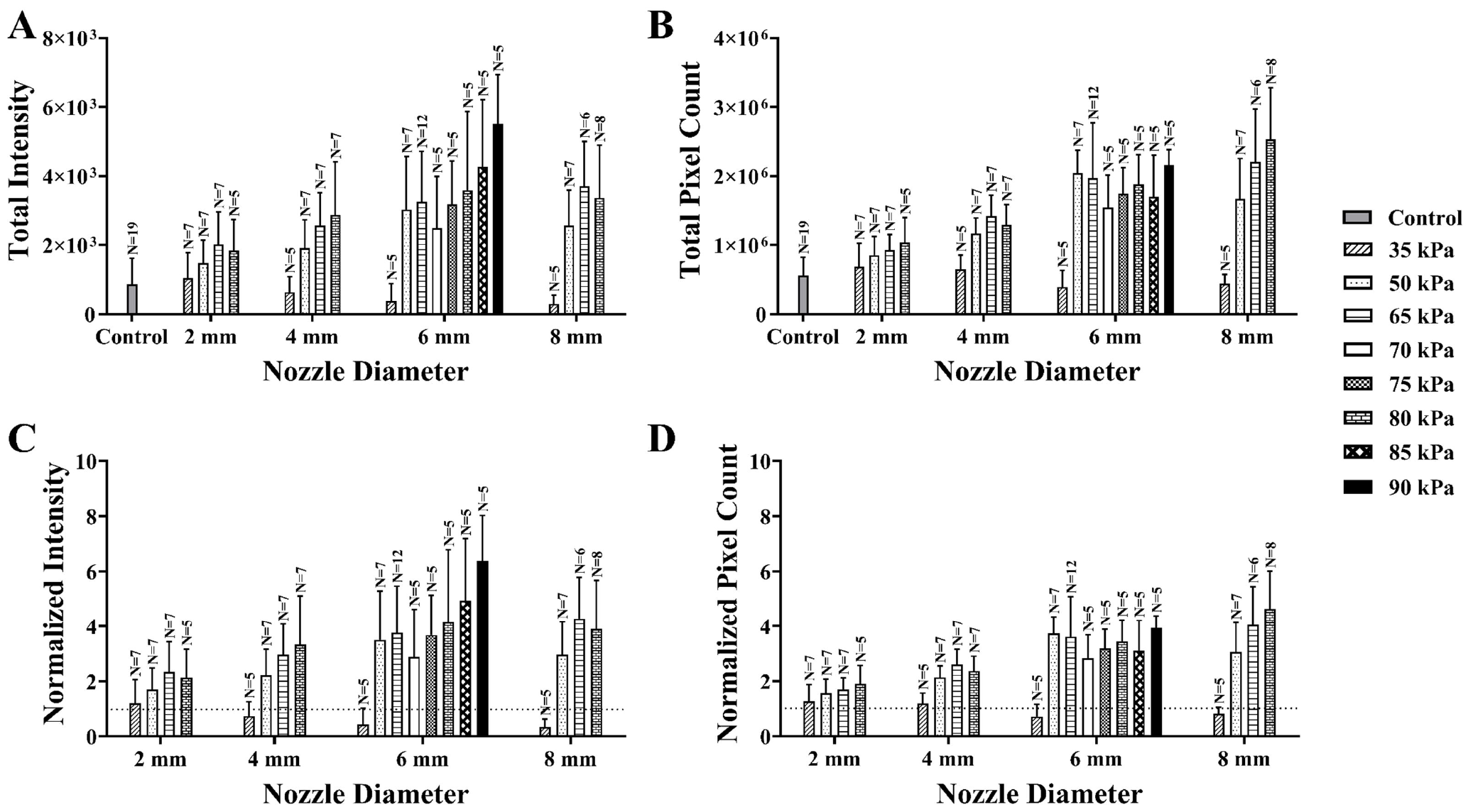
Variation of GFP expression with nozzle size and pressure. (A) Total intensity (mean+SD), (B) total pixel count (mean+SD), (C) normalized intensity (mean+SD), and (D) normalized pixel count (mean+SD) of GFP expression with different nozzle sizes and suction pressures. Note that normalized values represent fold-enhancement from injection-only control (dotted horizontal line). All suction pressures were applied for 30 seconds.

First, although expression increases in general with suction pressure for each nozzle size, the value of 50 kPa appears to be a threshold for significant enhancement. For the 6-mm nozzle size, no significant differences were observed among 50, 65, 70, and 75 kPa suction pressures. However, from 75 to 90 kPa, expression intensity levels are gradually increasing, and a positive correlation is observed between pressure and expression.

Second, with the exception of the lowest pressure (35 kPa), expression in general increases with nozzle size for the same applied suction pressure. (See also Fig. S1 where expression is shown as a function of nozzle size for each pressure.) For the 35 kPa group, no case resulted in expression enhancement above control, hence the positive correlation between nozzle size and expression is only valid from the threshold (~50 kPa) where expression enhancement is evident.

### Integrated total strain and tissue tension are indicators for effectiveness of suction-mediated transfection

To further analyze the strain-expression relationship, we performed a numerical simulation for each corresponding case and analyzed the resulting strain from suction-induced deformation. The integrated total strain (ITS) was computed as

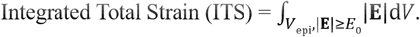

Here **E** is the strain tensor, and |**E**| is its Frobenius norm.^[13]^ V_epi_ is the volume of integration. In the depth direction, we integrate from the skin surface to 200 μm; this includes the epidermal layer, where we observed a majority of the expression.^[13]^ In the direction tangential to skin surface, we limit the integration to the area of whichever is smaller between the injection bleb and nozzle opening (see the schematic in Fig. S2). Furthermore, we impose a strain magnitude, *E*_0_, which we hypothesize represents a threshold for enhanced expression. The integration only includes those values above *E*_0_. Because this threshold is unknown, we varied *E*_0_ from 0-0.2, as presented in Fig. S3. The best correlation was achieved using a strain threshold, *E*_0_ = 0.15, which is shown in Fig. 5A and B. For the normalized intensity, two evident regimes were observed and are separated by the dotted line. The lower regime with normalized intensity below 1 is essentially where no-enhancement was observed; all 3 cases within this regime belong to the lowest pressure of 35 kPa. The upper regime exhibits a clear linear correlation between expression and ITS. In the upper regime, expression is correlated to ITS for both Normalized Intensity and Normalized Total Pixel Count, although the correlation is slightly stronger for intensity.

**Figure 5:**
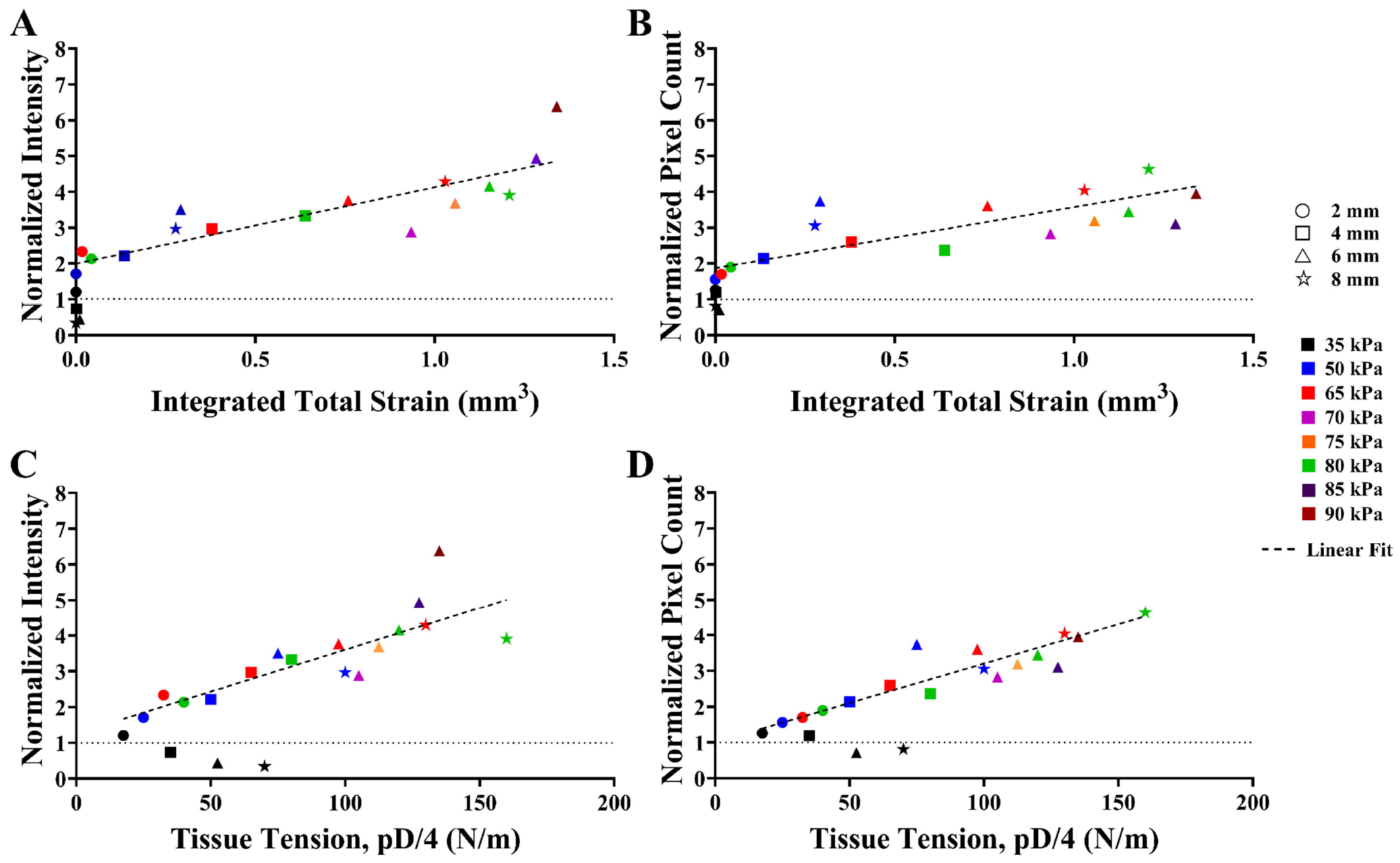
(A) Normalized intensity (mean) and (B) normalized pixel count (mean) of GFP expression as a function of total integrated strain, (C) normalized intensity (mean) and (D) normalized pixel count (mean) of GFP expression as a function of tissue tension.

Another useful and simpler indicator is the product of pressure and nozzle diameter (Fig. 5C and D). If the suction-induced dome is considered analogous to a liquid drop, this product is akin to the surface tension, and we include a factor of 1/4 to make the definition consistent with this parameter. The results again demonstrate that, excluding the same three cases for 35 kPa, a strong linearity appears between expression and tissue tension, rendering the latter an easy-to-use indicator to predict transfection efficacy using simple, circular-shaped nozzles.

## Discussion

In prior work, we have demonstrated that cutaneous, suction-based DNA delivery is an effective transfection method with efficiency comparable to other in vivo DNA delivery methods, such as electroporation and jet injection.^[13,33,42]^ This work builds on prior results and presents a systematic study of two of the most important parameters, nozzle size and pressure. Using the results of this study in combination with computational simulations, we have established a quantitative connection between strain and expression. In addition, we have also developed an alternative, simpler metric – tissue tension – that is well correlated to expression levels and does not require modeling for circular nozzles. The effectiveness of this indicator is not surprising as tissue tension is incorporated into the strain tensor. However, we have not yet identified a clear threshold for this metric, and as such it fails for the 3 cases with lowest expression (35 kPa: 4-, 6-, and 8-mm nozzle). Moreover, if more complex nozzle shapes are used, tissue tension is not as easily predicted, and modeling should be used.

We observed that the DNA cargo must be in the vicinity of cells during the application of suction to impart expression. This is both temporal – DNA injected immediately prior to suction generated strong expression, whereas DNA injected after suction had no impact; and spatial – we only observed expression in areas where DNA and suction were co-localized (Fig. 3). These results allude to important considerations in using suction as a means of enhancing expression following primary ID injection. Our prior study^[41]^ revealed that cargo for ID injection was contained within the visible bleb, especially for macromolecules such as DNA. Therefore, although in general tissue strain does scale with nozzle size for the same suction pressure, larger nozzles have limited benefits as they enclose areas greater than the bleb (see Fig. 3). In addition, because strain tends to concentrate toward the nozzle rim, a nozzle diameter approximately matching that of the bleb is preferred. For 50-μL ID injection, a 6-mm nozzle diameter provides the best matching. For the ID injection volumes of 100-200 μL (greater volumes are typically avoided because of leakage), the optimal nozzle dimeters would be 8 – 10 mm. The nozzle size should be matched to injection volume and bleb size to ensure co-localization and enhance expression.

The current and prior work indicate that a brief exposure to stress or strain leads to increased uptake of DNA. One mechanism for strain-induced uptake could be CLIC/GEEC (CG), a dynamin-independent endocytic pathway. In an in vitro study, Thottacherry et al. showed that activation of the CG pathway post-stretching is transient and vanishes as early as 90 seconds following strain-relaxation.^[43]^ Also consistent with the involvement of the CG pathway is the strain threshold required for triggering. We have demonstrated that using a strain threshold of 0.15 in evaluating total integrated strain reveals the best correlation with the observed GFP expression. This threshold value is commensurate with the several percent of linear strain needed to produce the necessary area dilation for CG-pathway activation;^[43]^ the details on this equivalence are elaborated in our prior work.^[13]^ To establish the CG endocytic pathway for suction-mediated transfection requires pharmacological studies using specific inhibitors (such as in ^[43]^ for in vitro fluid uptake) within the cutaneous DNA uptake context. However, it is evident that higher strain leads to higher transfection efficiency, and we anticipate that our current model can be an effective tool for optimization of nozzle geometric size and pressure selection.

## Conclusion

Application of suction to skin is a simple and effective method to enhance cutaneous nucleic-acid delivery and transfection. The current work presents a systematic study of delivery efficiency as a function of critical parameters, including nozzle diameter and suction pressure. We first demonstrated that suction needs to be applied after DNA injection, not before, to enhance transfection. We then performed a systematic study with various nozzle sizes and pressures, and used numerical modeling to correlate with GFP expression results. While expression depends on both parameters, the results indicate that expression level best scales with the total integrated strain above a threshold of 0.15. Furthermore, in the regime where suction-mediated transfection enhancement is evident, a simple variable, namely, the estimated tissue tension, can be used to predict expression level. These results provide useful relationships and guidelines toward the optimization of this platform. Future work includes pharmacological studies to confirm the endocytic pathways responsible for cellular uptake. Although currently we focus on cutaneous suction and uptake, we anticipate that the platform can be extended to other organ/tissue applications.

## Data Availability Statement

The data of this study is included in the article and supporting information. Further queries can be directed to the corresponding authors.

## Ethics Approval Statement

All animal experiments were approved by Rutgers University Institutional Animal Care and Use Committee (IACUC), protocol: 201800077.

