## Supplemental Materials for "Tissue Tension and Strain as Indicators of Suction-mediated Cutaneous DNA Transfection: A Parametric Study"

### Title


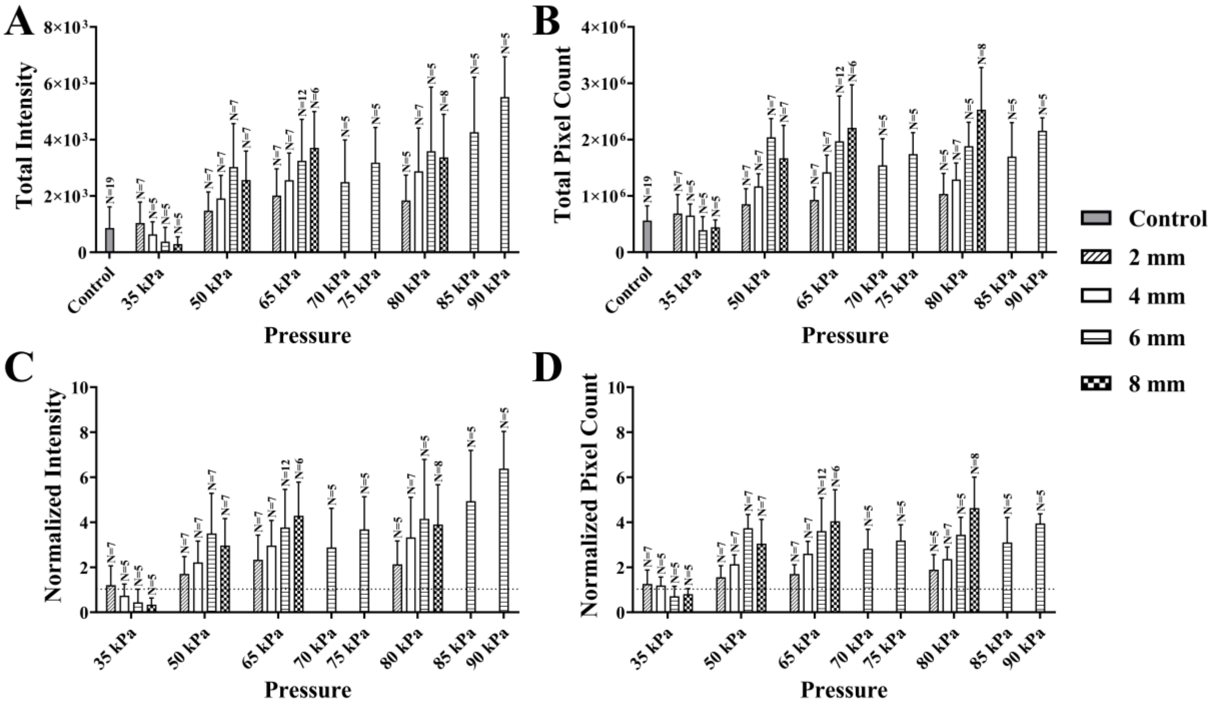


**Figure S1:** Variation of GFP expression with nozzle size and pressure. Here the intensity (mean±SD) (A), (C) and pixel count (mean±SD) (B), (D) of GFP expression are grouped with respect to suction pressure. All suction pressures were applied for 30 seconds.


†
‡

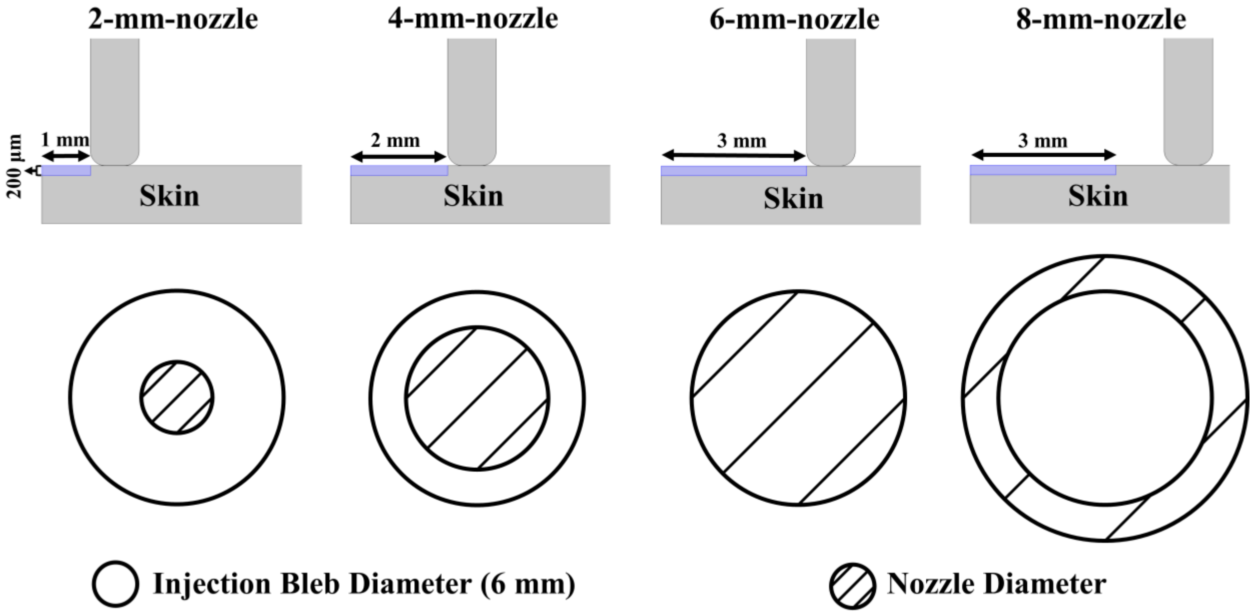


**Figure S2:** Determination of $V_{epi}$ , indicated by the blue shaded area. The volume of integration is determined by considering both cargo distribution and axisymmetric nozzle size. For nozzle sizes smaller or comparable to injection bleb, namely, D=2, 4, and 6 mm, the area underneath the nozzle opening is used. For nozzle size greater than the bleb, namely, D=8 mm, the bleb size is used. See also Fig. 3 in the proper text for reference.

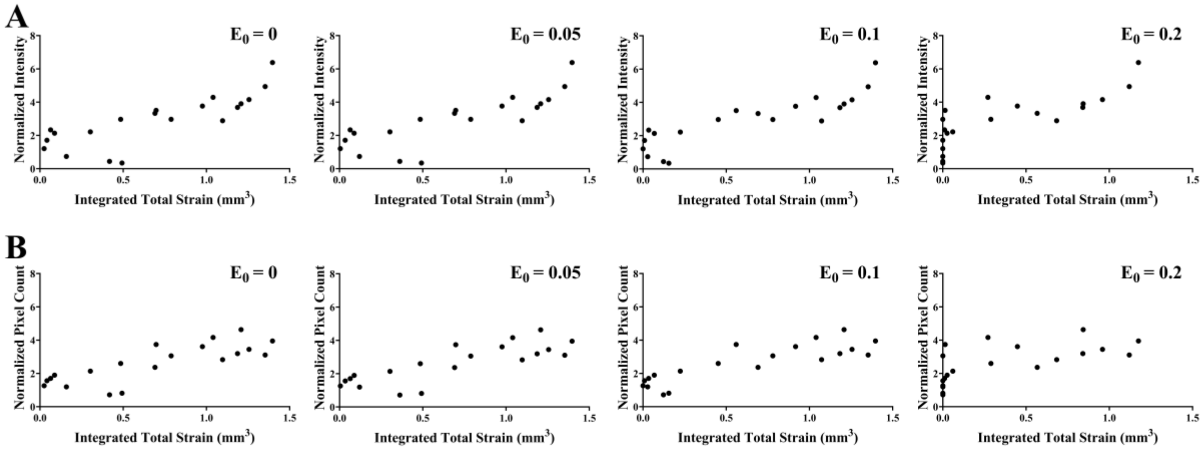


**Figure S3:** (A) Normalized intensity (mean) and (B) normalized pixel count (mean) of GFP expression as a function of total integrated strain where different strain thresholds of $E_0$ are chosen; the case of $E_0$=0.15 is shown in Fig. 5 in the proper text.
